# Time Perception through the Processing of Verb Tenses: An ERP study regarding Mental Time Travel

**DOI:** 10.1101/2020.12.23.424164

**Authors:** Charalabos Papageorgiou, Anastasios E. Giannopoulos, Athanasios S. Fokas, Paul M. Thompson, Nikolaos C. Kapsalis, Panos Papageorgiou, Xanthi Stachtea, Christos N. Capsalis

**Author notes:** Correspondence: Athanasios S. Fokas, and Anastasios E. Giannopoulos.

## Abstract

Humans are equipped with the so-called Mental Time Travel (MTT) ability, which allows them to consciously construct and elaborate past or future scenes. The mechanisms underlying MTT remain elusive. This study focused on the late positive potential (LPP) and alpha oscillations, considering that LPP covaries with the temporal continuity whereas the alpha oscillations index the temporal organization of perception. To that end, subjects were asked to focus on performing two mental functions engaging working memory, which involved mental self-projection into either the present-past (PP) border or the present-future (PF) border. To evaluate underlying mechanisms, the evoked frontal late positive potentials (LPP) as well as their cortical sources were analyzed via the standardized low-resolution brain electromagnetic tomography (sLORETA) technique. The LPP amplitudes - in the left lateral prefrontal areas that were elicited during PF tasks - were significantly higher than those associated with PP, whereas opposite patterns were observed in the central and right prefrontal areas. Crucially, the LPP activations of both the PP and PF self-projections overlapped with the brain’s default mode network and related interacting areas. Finally, we found enhanced alpha-related activation with respect to PP in comparison to PF, predominantly over the right hemisphere central brain regions (specifically, the precentral gyrus). These findings confirm that the two types of self-projection, as reflected by the frontally-distributed LPP, share common cortical resources that recruit different brain regions in a balanced way. This balanced distribution of brain activation might signify that biological time tends to behave in a homeostatic way.

## INTRODUCTION

The so-called Mental Time Travel (MTT) is an important mental skill that results from our capacity to be aware of subjective time. MTT enables us to re-experience past events and to imagine possible future events (Tulving,1985).

Modern debates regarding the origin of MTT are strongly influenced by the crucial relationship of MTT with language (Suddendorf and Corballis, 2007). Language, among other things, reflects the structure of the perception of time, as shown by the development of verb tenses (“X” happened yesterday, “Y” is happening now, “Z” will happen tomorrow). In this sense, it is reasonable to assume that the structure of time is reflected in the structure of grammar. This relationship was clearly understood by the Greek sophist Protagoras (490 – 420 BC), who was the first to distinguish the tenses and to emphasize the importance of the movement of time (Dillon and Gergel, 2003). In principle, the mental representation of time is commonly determined through descriptive terms, such as the “arrow of time” or “time passage”; these are attempts to assign time characterizations of past, present, and future events. Time perception is connected with the deep intuition that the future can be changed until it become s present, but the past is fixed. In other words, events that have not occurred are potentially alterable, unlike the unaltered events that have already happened (McCormack, 2014). Constructing a future event with all its ‘unseen’ details, in general requires greater mental effort than recalling a past event. This structure of the “fixed past”, “immediate present” and “open future” is deeply engrained in our language, and in our thoughts and behavior (Callender, 2010; Oppenheim, 2010). Philosophically, this concept was elaborated at the beginning of the last century by philosopher John McTaggart in his A-theory (or tensed theory) of time (McTaggart, 1908). According to this theory, all events are characterized in terms of their temporal specification, namely as being past, present, or future (Oaklander, 1996). We perceive events (instances in time) approaching from the future, passing by in the present, and receding into the past (time-moving metaphor); also, we perceive objects (including our sense of self) travelling through time from past to future (ego-moving metaphor) (McGlone and Harding, 1998; Thönes and Stocker, 2019).

Behavioral and brain imaging studies show robust mental signatures of MTT in humans and in animals (Suddendorf and Busby, 2003; Viard et al., 2011). Several studies have shown that the dorsolateral pre-frontal cortex (DLPFC) mediates the working memory aspects of the timing, whereas the activation of the prefrontal, premotor, and anterior cingulate cortices, is related to attentional aspects of time perception (Fuster, 2015; Fuster and Bressler, 2012; Lewis and Miall, 2006; Nobre and O’Reilly, 2004). Experiments requiring mental time projection typically engage working memory (WM) operations. WM is believed to be a system for temporarily storing and managing the information required to carry out complex cognitive operations, including reasoning (Baddeley, 2012; Hasson et al., 2015).

Other research suggests that the late positive component (LPC) - or late positive potential (LPP) - of event-related brain potentials (ERPs) may reflect successful decision-making regarding time estimation, as a result of neuronal activity in the PFC (Gontier et al., 2009; Paul et al., 2003, 2011). Recently, additional evidence showed that LPP amplitude covaries with the difficulty of temporal discrimination of continuity (Wiener and Thompson, 2015). By using the term *temporal continuity,* we refer to the “aspect of conscious perception that moments carry over from one to the next” (Wiener and Thompson, 2015).

With regard to the origin of the LPP, LPPs may reflect both prefrontal activity induced by attention-arresting stimuli, and the mental requirements of working memory (Gibbons et al., 2018; Kopf et al., 2013). Moreover, recent studies propose that alpha brain oscillations may reflect the temporal sequence coordination of the associated neuronal representations. The underlying rhythmic neuronal oscillations operate as an attentional ‘gatekeeper’ that allocates priority to certain stimuli for WM storage by enabling an optimal signal-to-noise ratio; in this way, possible interference with conflicting sensory inputs (Freunberger et al., 2011; Grabot and Kayser, 2020; Jensen et al., 2014) is avoided or reduced. In this framework, a considerable body of evidence has highlighted a relationship between the alpha phase and timing in perception. Specifically, there may be a distinct role for the alpha oscillatory activity in determining temporal resolution (Milton and Pleydell-Pearce, 2016; Samaha and Postle, 2015; VanRullen, 2016). Notably, Ronconi et al., (2017) demonstrated that alpha EEG oscillations provide a hierarchical framework for the temporal organization of perception.

Building on these foundations, here we attempted to integrate two key research directions – namely, LPP and alpha EEG brain oscillations evoked by the two diverse forms of *self-projection* in time. Noteworthy, self-projection refers to the mental ability to shift our perspective from the immediate present to alternative past or future perspectives of a certain event. In this study, triplets of verb tenses (past, present, future) were used to enable the self-projection in time, triggering the perceptual shift from the present tense of a verb towards its alternative - past or future - tenses. By analyzing evoked LPPs, we evaluated frontal brain activation elicited during the processing of tensed verbs (based on the ‘A-theory’ by McTaggart, 1908). Specifically, participants were asked to project themselves either into Past-Present (PP) borders (i.e., “from near past to present”) or into Present-Future (PF) borders (i.e., “from present to near future”). Phrases in parentheses were borrowed from McTaggart’s terminology (McTaggart, 1908) (see **Appendix** for more information on the “borders” terminology).

Scalp evoked LPPs were processed using the standardized low-resolution electromagnetic tomography (sLORETA) technique (Pascual-Marqui, 2002). This approach provides estimates for the cortical distribution of the electrical ERP generators, and allows us to differentiate the source-level resources between the two experimental tasks (PP vs PF).

Recently, our group developed a novel method, which solves an inverse problem to determine a near- to far-field transformation of the source EEG data; using this approach, we found that the overall power emitted by a single subject undergoing a self-projection into the PF borders, was greater than that corresponding to the PP self-projection. These findings have been discussed in relation to the biological effect of the second law of thermodynamics (Katsouris et al., 2019). This is consistent with the existence of a direct relationship between the passing of time and increased entropy (Ghaderi, 2019).

Based on the above considerations, several hypotheses were put forward. First, we predicted that the frontally-distributed LPP, observed in our study, would provide a useful dissociative tool to elucidate the role of the PFC network. As time estimation engages MTT ability as well as WM, we hypothesized that the above network would be involved in the processing of tensed tasks (verbs). Second, based on the biological realization of the second law of thermodynamics - and the fact that future thinking involves a process of actively constructing unknown events - we hypothesized that the measures of brain-derived signal entropy, as reflected by the neuronal sLORETA activations, would be enhanced during the future-related (PF) task. Third, considering the role of alpha activity with respect tothe temporal coding organization in the brain, we predicted that the alpha oscillatory activity would reflect complementary mechanisms influencing self-projection intoboth the past and future.

## MATERIALS AND METHODS

### Participants

Thirty-nine healthy adults (mean age: 25.3 y ± 2.8 SD; 15 males; 35 right-handed; educational level: 16.9 y ± 0.9 SD) participated in the experiment. All participants were volunteers with no history of mental illness or brain disease. All gave written consent, after being extensively informed about the procedure. Inclusion criteria for all participants were the absence of medical, neurological, psychological problems, and any pharmacological treatment. Before starting the recording session, each participant was trained with 3 trials to familiarize themselves with the experimental material. Participants had no previous experience of the aims of the study. The “Protagoras” experiment (Katsouris et al., 2019) described below was carried out in accordance with the Code of Ethics of the World Medical Association (Declaration of Helsinki) for experiments involving human participants.

### “Protagoras” Experiment: Procedure and Stimuli

The experimental procedure was designed based on a two-tone paradigm. Participants sat in a comfortable chair and were instructed that they would hear two successive tones (“beeps”) per trial through headphones. Before the first tone onset, a *verb triplet* was heard through the headphones with a speech intensity level of 65 dB. Verb triplet refers to the same verb in its three tenses - the past, present, and future tense (e.g., for the verb “love”, the triplet is “I loved, I love, I will love”).

During the fore period (time interval between the two tones), the subject, depending on the tone frequency, mentally concentrates on either the past/present or present/future tenses of the verb triplet. If the tone frequency is 3 kHz the concentration target is the past-present tense; if the tone frequency is 500 Hz, the concentration target is the present-future tense. The single-trial recording is completed when the second tone is triggered. Before the next trial recording begins, the subject has to declare a degree of confidence about its task-relevant concentration level (from 0% corresponding to zero concentration, to 100% associated with perfect concentration). Both tones had a duration of 100ms, same frequencies (both at 3 kHz or both at 500 Hz) and were separated with an inter-stimulus interval (ISI) of 3 seconds. The inter-trial intervals varied in duration from 4 to 9 seconds. A single-trial recording is graphically presented in **Fig. 1**. The fore period is considered necessary to elicit EEG and ERP factors that uncover how information is processed and to influence the response preparation.

**Figure 1.**
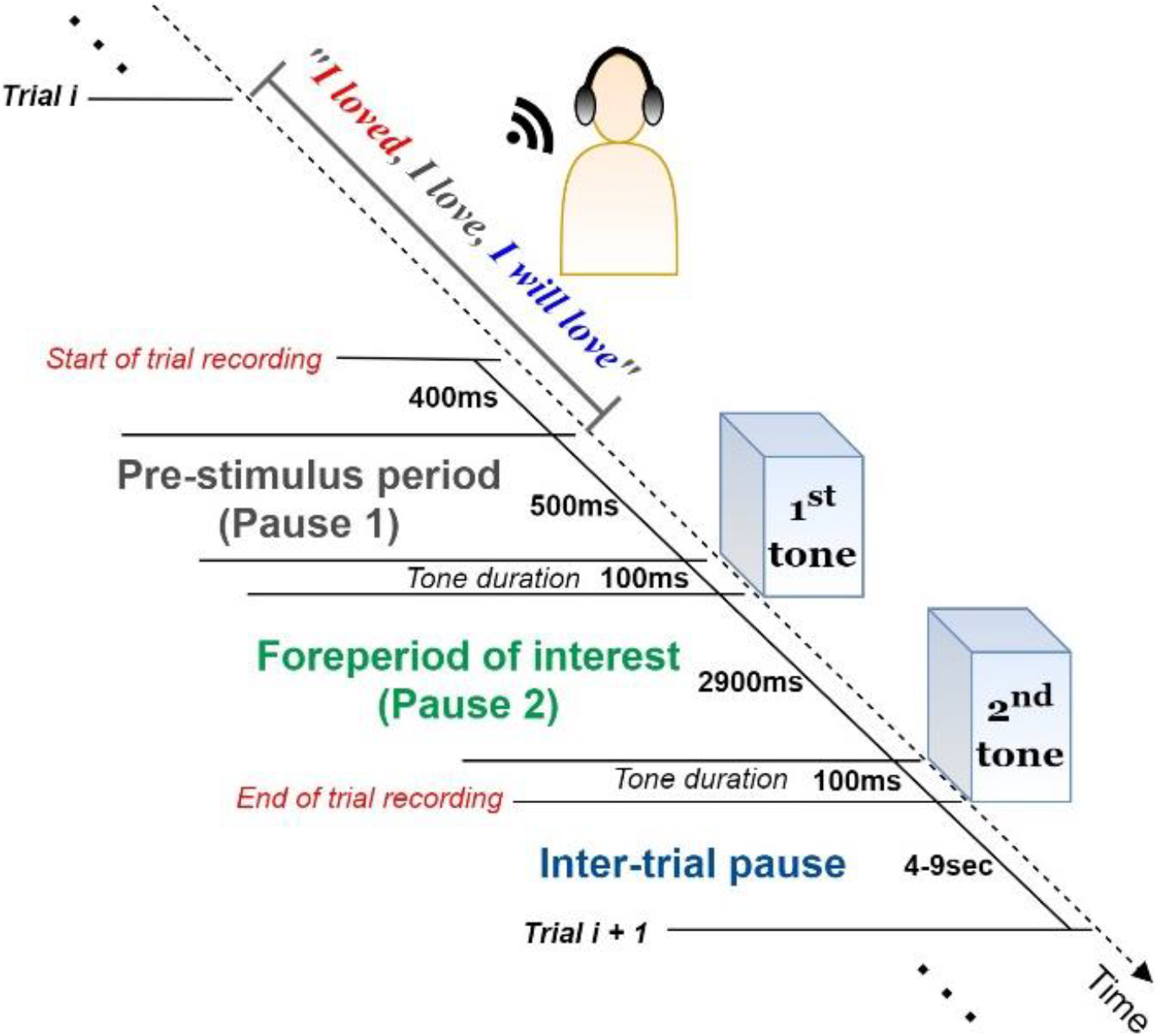
Single-trial structure. (a) The subject listens carefully to the verb triplet through a pair of earphones. (b) EEG pre-stimulus activity is recorded for a time period of 900 ms before the target stimulus(S1) arousal. (c) Depending on the frequency of S1, the subject is asked to concentrate on the past/present (if S1 is at 3 kHz) or present/future (if S1 is at 500 Hz) tenses of the verb. EEG post-stimulus activity is captured for a time period of 2900 ms until the second stimulus (S2) arousal. (d) During the between-trials pause (4-9 sec), the subject declares their degree of confidence (%) regarding their concentration performance on the particular verb.

Moreover, these factors are highly informative about the temporal order of information processing and the task-related performance of the subject’s brain.

The above recording structure is repeated 210 times during a session (105 different verbs × 2 different target tenses). The same verb triplet is presented twice during a session: once for concentrating on the past/present and once for concentrating on the present/future tenses. Therefore, each participant is tested for a total of 210 trials, under two experimental conditions: 105 trials targeting the past tense and 105 targeting the future tense. To prevent habituation effects, the order of the verbs was pseudorandom across participants; to avoid tiredness,the between-trials interval (4 up to 9 seconds) is manually chosen by the subjects.

### Data Acquisition

EEG recordings were conducted in a Faraday cage to minimize interference from external electromagnetic fields. A Line Impedance Stabilization Network (LISN) was used to eliminate possible conducted emissions. Evoked biopotential activity was digitalized at a sampling frequency of 1 kHz (sampling period 1 ms) from 30 scalp sites (FP1, F3, P3, O1, F7, T3, T5, AFz, Fz, FCz, CP3, FC3, TP7, FPz, FT7, Oz, FT8, FP2, F4, C4, P4, O2, F8, T4, T6, Cz, Pz, CPz, CP4, FC4) using active electrodes mounted on an elastic cap, in accordance with the International 10-20 System. To detect blinks or eye movements, horizontal (HEOGs) and vertical (VEOGs) electro-oculograms were recorded from two electrodes. The VEOG electrode was placed above the right eye, whereas the HEOG electrode was placed at the outer canthi of the left eye. Electrode impedance was kept constantly below 5kΩ. The brain signals were amplified (gain 47 dB) by a Braintronics DIFF/ISO-1032 amplifier before entering a 32-bit analogue to digital converter (NI SCB-68), which has a GPIB output. The digitized signal comprised an input for National Instruments PCI-6255 DAQ card (16 bits ADC) through two National Instruments CB-68LP terminal blocks. The PC with the DAQ Card runs a LabView program for the recording of the signals, which can be monitored by an on-screen graphical representation. EEG online activity was referenced to the ear lobes while the ground electrode was placed on the left mastoid.

### Preprocessing pipeline

All datasets were preprocessed using the EEGLAB environment and denoising functions (Delorme and Makeig, 2004). Firstly, EEGs were down-sampled to 250 Hz, to compress the data size and suppress unnecessary high-frequency information for the Independent Component Analysis (ICA) decomposition. The data were then band-pass filtered (using the default FIR filter of EEGLAB) in the band 0.1-45 Hz to remove the baseline “drifts” and ignore the 50-Hz line noise. Using the “clean_rawdata” function, an EEGLAB plugin for bad channel detection (see EEGLAB documentation), electrodes showing abnormal time-course were excluded (no more than three channels per participant). After the interpolation of all removed channels, each electrode activity was re-referenced to the whole-scalp common average. Electrodes FP1, FP2 and FPz were not considered reliable due to their noisy time-course (they were replaced by interpolation in 28 of the single - subject datasets).

To eliminate the contribution of non-brain components (especially blinking and saccades) from the measured data, the resulting datasets were decomposed via the ICA algorithm (Delorme et al., 2007), providing estimates of independent component (IC) activations. Furthermore,the SASICA tool provided by EEGLAB plugins was employed to guide the selection of non-brain components (Chaumon et al., 2015). Artefactual components removal was performed semi-automatically, including visual inspection of IC time-course, spectra and topography, along with simultaneous consideration of the SASICA guidelines parameterized via: “Autocorrelation” (Threshold (r) = auto; Lag = 20ms), “Focal components”(Threshold (z) = auto), “Correlation with EOG” (enabled for VEOG and HEOG with threshold (r)= 0.2), “ADJUST” (Mognon et al., 2011) and “FASTER” (Nolan et al., 2010) methods (enabled for blink channels). Finally, artifact-free data were obtained by reconstructing the remaining non-artifactual ICs in the scalp domain. Before further processing, the continuous data were segmented into 2.5 -second epochs (−0.5 to +2 sec), they were time-locked to the first tone onset, and were baseline-corrected based on the 200-ms pre-tone period.

Trials corresponding to the zero declared degree of concentration confidence (0%), were excluded from subsequent analyses. There were no significant differences in trial count across participants (maximum number of zero-concentration trials per subject was three).

### Event-Related Potentials

Scalp-domain ERP waveforms were extracted by separately averaging the same-condition trials (2 electrode-specific ERPs per subject). To visually inspect the “when-and-where” of LPP elicitation, and to obtain a “general task engagement” view of the ERP waves, the grand-averaged ERPs (across participants and conditions) were calculated in each electrode.

Based on visually inspecting the grand-averaged scalp maps, a broad (~400-800 ms) anterior positive deflection (peaking within 500-600 ms) was observed in both conditions (see **Fig. 2**). To reduce the spatial dimensionality of the analysis, we analyzed three frontal regions of interest (ROIs): left centro-frontal sites (LFC; F7, FT7, F 3, FC3), right centro-frontal sites (RFC; F8, FT8, F4,FC4) and midline centro-frontal sites (MFC; AFz, Fz, FCz). Each of these ROIs was represented by the average ERP wave across its electrodes, thus minimizing the familywise error rate in the statistical testing (Luck, 2014). We used this ROI approach (rather than analyzing each single frontal electrode) to reduce the number of comparisons, avoid noisy (close to the eyes) channels (FPz, FP1 and FP2), examine lateralized (left, right, midline) effects and focus only on the areas where LPPs are positive-and-maximal, as they were observed in collapsed (across participants and conditions) ERPs. The LPP component was analyzed for each single-subject ERPs in the post-stimulus window of 400-800 ms. Given the well-documented robustness and high signal-to-noise ratio properties of mean measures against peak detections (Luck, 2014), single-subject LPP amplitudes were extracted as mean values within the above window.

**Figure 2.**
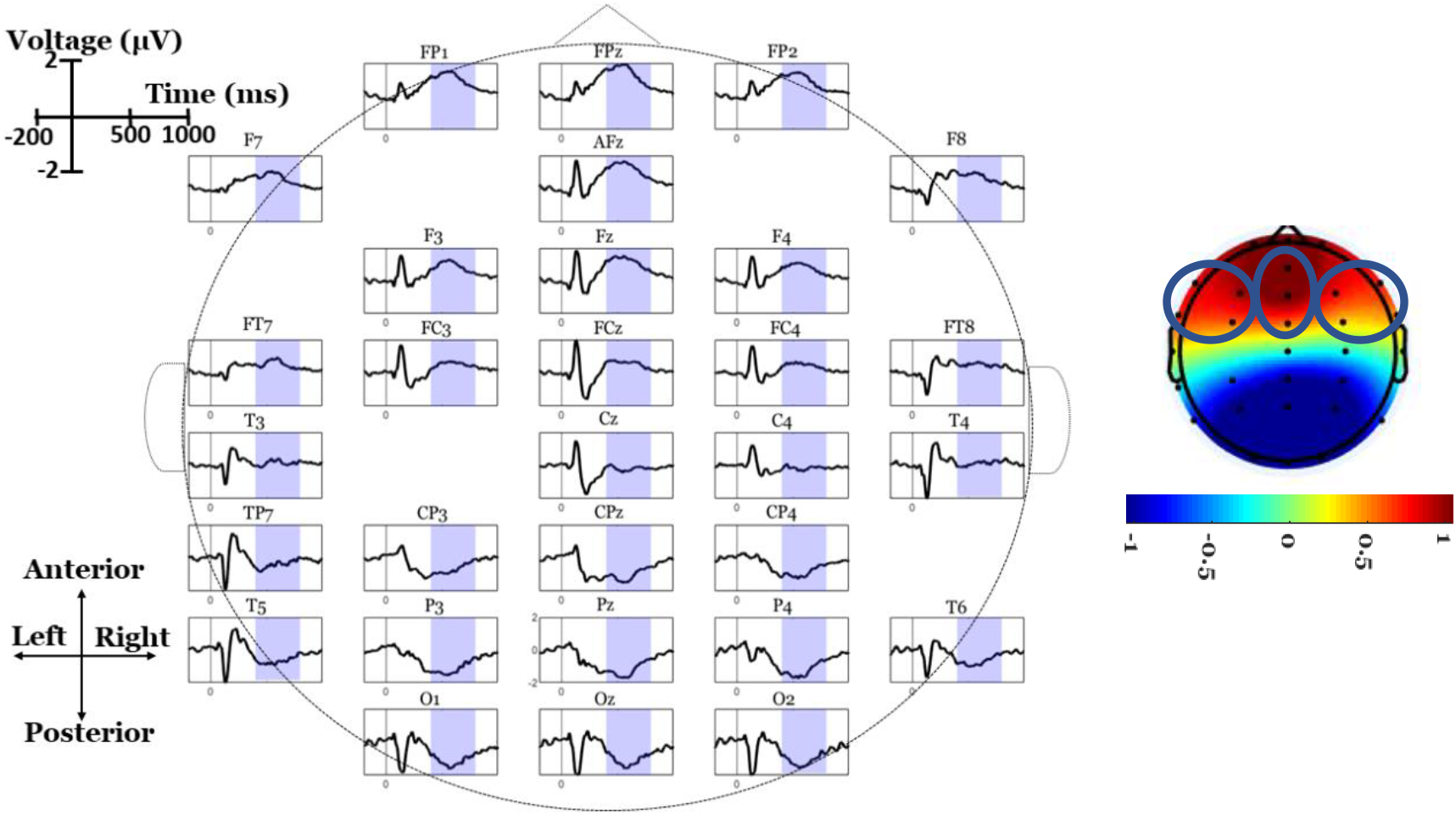
Grand-average ERP waves (across participants and conditions) at each electrode. Blue-shaded areas indicate the LPP window (400-800 ms). On the upper right, mean scalp topography is illustrated (average voltage across 400-800 ms) for LPP, as well as the three ROIs.

### Identifying ERP brain sources with sLORETA

ERP responses were exported for further analysis using the sLORETA software. In general, the sLORETA inverse-problem solution algorithm has been established as a reliable estimator of (sub)cortical sources, providing a useful approach for the analysis of different time segments of ERPs (Decety et al., 2010; Nir et al., 2008; Schneider et al., 2009). The sLORETA solution space detects source localizations in 6, 239 cortical gray matter voxels with a spatial resolution of 5×5×5 mm; localization inference is based on standardized values for current density estimates (Wagner et al., 2004). The implementation incorporates a 3-shell spherical headmodel registered to a recognized anatomical brain atlas (Talaraich and Tournoux, 1988). sLORETA enables the computation of statistical maps from ERP components data that indicate the locations of underlying generators with low error (Pascual-Marqui,2002).

First, the 30 electrode coordinates were positioned using the Talairach coordinate system according to the spatial association between anatomical brain landmarks and scalp positions (Towle et al., 1993). These Talairach coordinates were then used to compute the sLORETA transformation matrix. Using the transformation matrix (without any smoothing), condition-specific ERPs for each subject were transformed to sLORETA files, containing the 3D cortical current source density vectors (magnitudes) of each voxel. Finally, the source localization of LPP was calculated as the mean sLORETA image (mean activations within 400-800 ms).

### Identifying band-specific brain sources with sLORETA

Apart from the time-domain source localizations,sLORETA was used to estimate band-specific sources within the delta (1-4Hz), theta (4-7Hz), alpha (8-13Hz), beta1 (16-24Hz) and beta2 (25-30Hz) bands. For this purpose, ERP data were fed into the sLORETA module extracting the scalp-domain cross-spectra measures related to LPP evoked oscillations. Cross-spectra files were finally transformed into the 6,239-source domain providing estimates of the LPP band-specific voxel activations.

Based on EEG data, it is not possible to re construct the neuronal current uniquely (Dassios and Fokas, 2020). A novel algorithm toreconstruct the ‘visible’ by EEG part of the current, namely the part of the current that affects the EEG data, is presented in a recent study (Hashemzadeh et al., 2020). This algorithm uses real brain topology without the spherical approximation. Due to certain technical difficulties, it was not possible to use the latter algorithm in the present work; however, there is a broad agreement between the results obtained via the LORETA technique and the approach of this algorithm (Hashemzadeh et al., 2020).

### Statistical analysis of scalp differences

To detect statistically significant effects in the scalp data, a 2-by-3 repeated-measures analysis of variance (ANOVA) was conducted on LPP amplitudes. The ANOVA factors were Condition (PP vs PF) and ROI (LFC vs RFC vs MFC). Interaction effects were addressed by juxtaposing PP and PF conditions in each ROI using paired *t*-tests. To adjust for multiple statistical comparisons, all *post hoc* tests used the Bonferroni-adjusted *p* = .05/3 = .0167.

### Statistical mapping of brain activations

The sLORETA software was also used to statistically map 3D cortical distribution differences using a non-parametric approach (Nichols and Holmes, 2002). First, condition-specific LPP images are compared against baseline (−200-0ms) looking for voxels exceeded the mean baseline value by at least 3 standard deviations. This was performed by replacing each baseline value with *Baseline*_*mean*_ + 3 × *Baseline*_*STD*_ (where STD stands for “standard deviation” of the mean), and then contrasting LPP activations against the (modified) baseline. This procedure detected LPP-related sources against pre-stimulus (background) sources. In addition, LPP images were contrasted between the PP and the PF conditions, to identify voxels that were more active in the PP versus PF condition or *vice versa*. Regarding the band-specific source comparisons, we performed only contrasts between conditions considering the LPP activations as the dependent variable. All statistical thresholds were set to the critical *t*-value (log-of-F-ratios option of sLORETA) corresponding top<.05, as defined by 5,000 randomizations (Nichols and Holmes, 2002); the results were displayed as *t*-statistic brain maps.

## RESULTS

### Behavioral measures

At the end of each trial, participants stated their degree of concentration in the given task. The mean concentration score for each condition was calculated per participant. To test whether the reported degree of concentration is affected by the experimental condition, we compared the mean concentration levels between PP and PF tasks using a paired *t*-test (two-tailed, critical *t*-statistic = ±2.024). Reported concentration scores were significantly higher (t(38)=2.65, p=.012) in PP (67.98 ± 2.31%) than PF (65.81 ± 2.52%) trials.

### LPP scalp differences

The mean amplitudes of frontal LPPs were tested for possible alternations between PP and PF conditions. ANOVA testing revealed a significant interaction effect 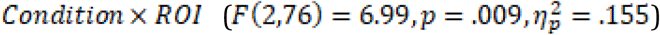. The condition-specific LPPs were then contrasted in each ROI, separately, showing that LFC areas measured higher amplitudes in PF than PP condition (*t*(38) = −2.55, *p* = .015), whereas the opposite pattern was observed for the MFC (*t*(38) = −2.59, *p* = .013) and RFC (*t*(38) = 3.27, *p* = .002) sites. The grand average ERPs, scalp topographies and descriptive statistics are illustrated in **Fig. 3**.

**Figure 3.**
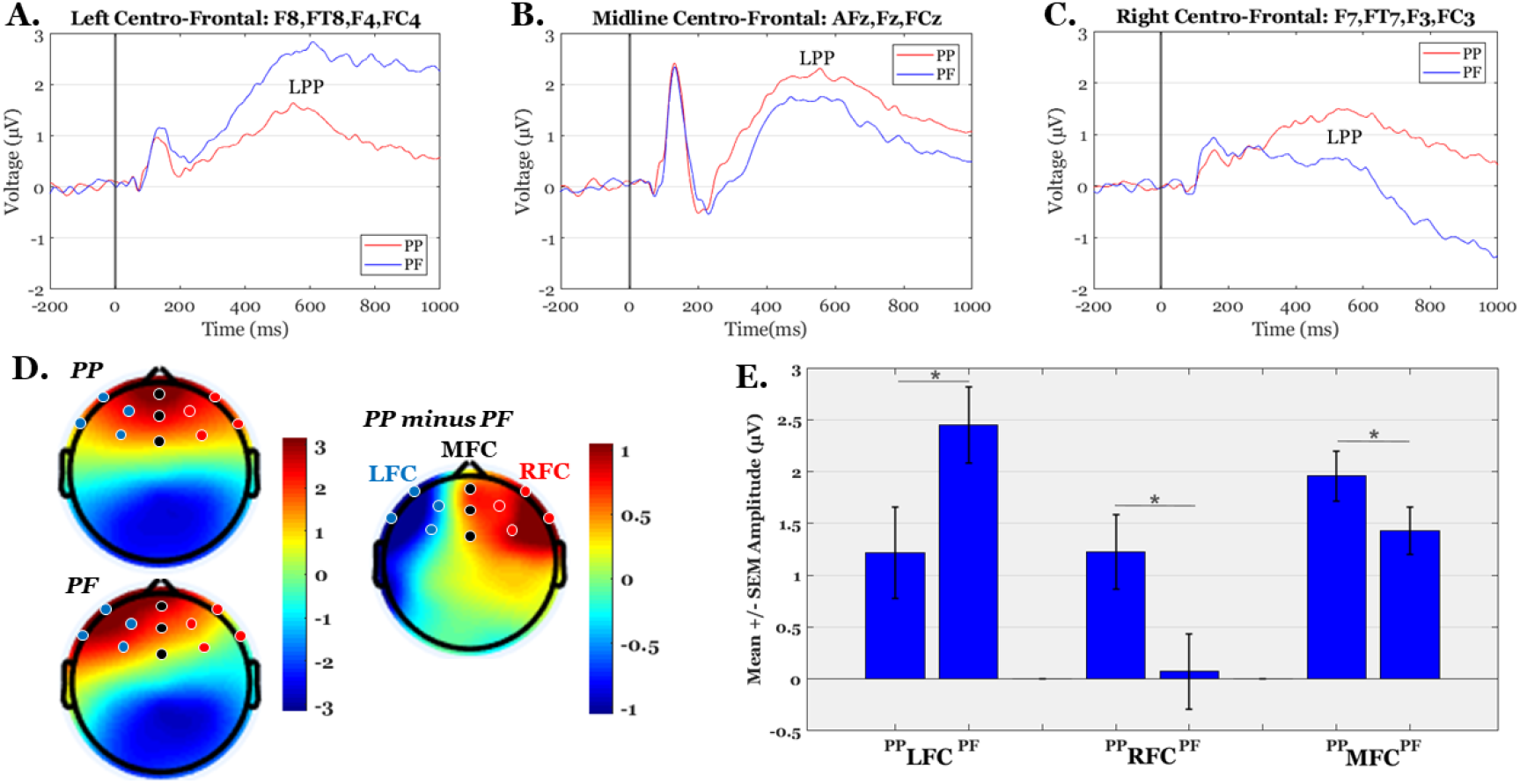
Grand-average ERP waves, scalp maps and descriptive statistics of LPP amplitudes. Panels A-C illustrate the grand-average ERPs over Left-Fronto-Central (LFC), Midline-Fronto-Central (MFC) and Right-Fronto-Central (RFC) regions, respectively. Panel D shows the mean scalp topographies of LPP (average within 400-800 ms) in PP, PF and their difference (PP-PF). Panel E shows the Mean +/− SEM (Standard Error of the Mean) of LPP amplitudes in PP and PF conditions over the three ROIs (‘*’ indicates significant differences at p<.05).

The behavioral data (concentration scores) and electrophysiological data (LPP amplitudes) were also tested for possible correlations (using matrices of Pearson’s correlation coefficients). No significant relationships were detected between any pair of ROI-specific LPPs and concentration scores (all *p*’s>.20).

### Source localization of the LPPs

Source localization of the entire scalp topography during the time frame of LPPs, revealed significant activations in several brain regions; these areas are tabulated in **Table 1** (for the PP condition) and **Table 2** (for the PF condition). We reported the (voxel) clusters for which at least five significant voxels adjacent in 3D space (significance threshold of *p*<.05; critical *t* is reported in the color bars). Both conditions revealed the highest activations around the anterior cingulate regions (BA 33 for PP; BA 24 for PF). **Figures 4A** and **4B** demonstrate the LORETA images for the PP and the PF sources, respectively, in *xyz*-slices that correspond to the maximum-*t* views. For completeness and comparison purposes, **Figure 4C** shows the PP and PF sources in the same plot from six different 3D views.

**Table 1.**
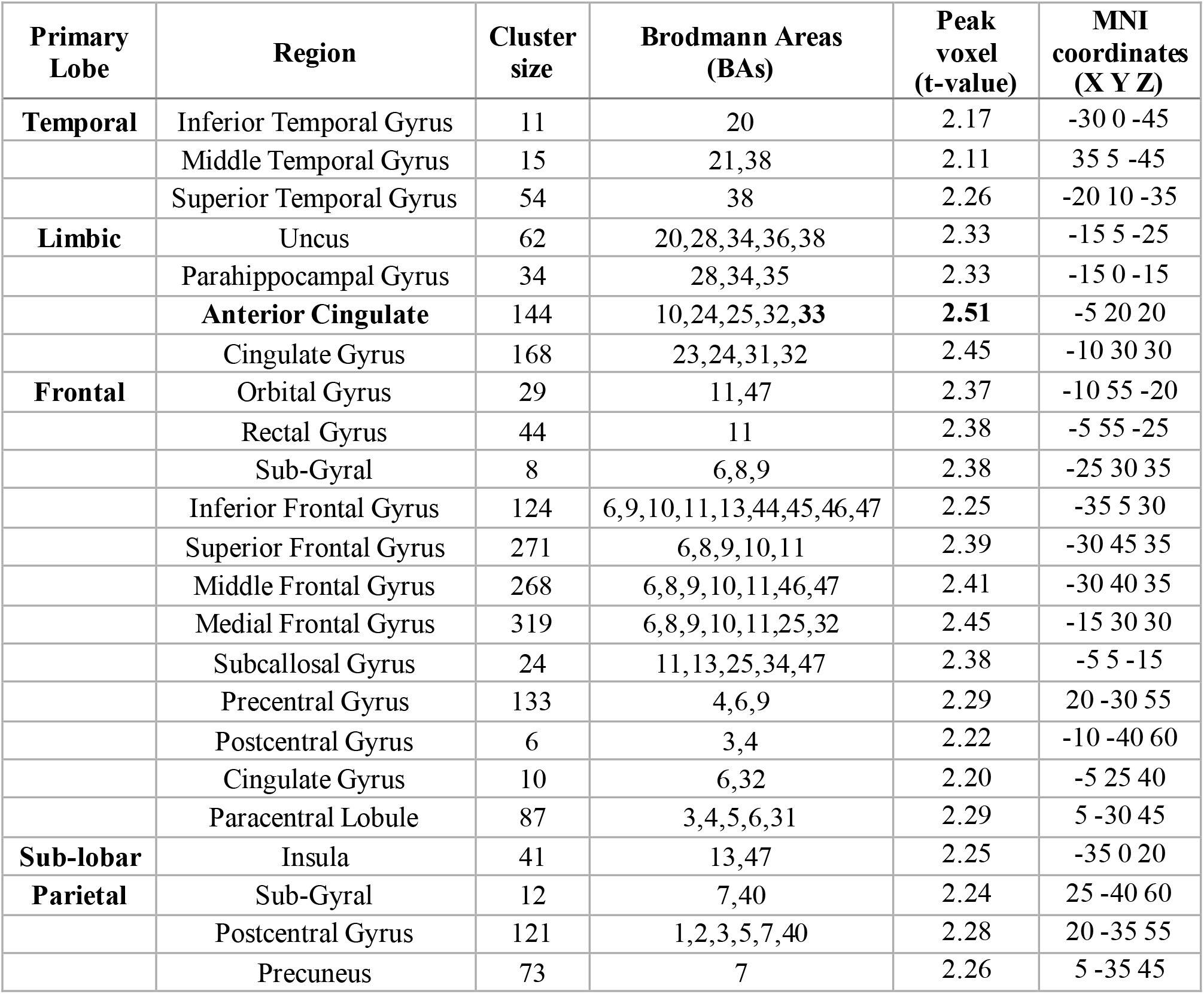
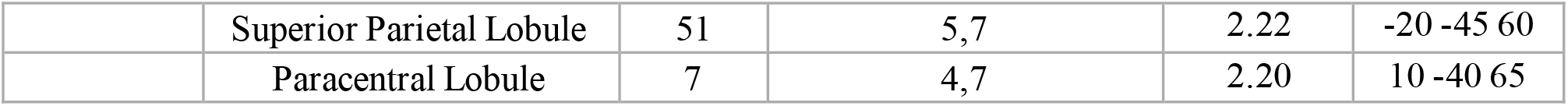
Localization of scalp sources’ response to the PP during the time window characterizing LPPs. Elementsin **bold font** indicatethe maximal t-scores.

**Table 2.** Localization of scalp-sources’ response to the PF during the time window characterizing LPPs. Elementsin bold font indicate the maximal t-scores.

| Primary Lobe | Region | Cluster size | Brodmann Areas (BAs) | Peak voxel (t-values) | MNI Coordinates (X Y Z) |
| --- | --- | --- | --- | --- | --- |
| **Temporal** | Middle Temporal Gyrus | 5 | 21,38 | 2.06 | 35 10 -45 |
|  | Superior Temporal Gyrus | 32 | 38 | 2.15 | 25 10 -30 |
| **Limbic** | Uncus | 41 | 20,28,34,36,38 | 2.18 | 20 5 -30 |
|  | Parahippocampal Gyrus | 15 | 28,34,35 | 2.17 | 20 5 -20 |
|  | **Anterior Cingulate** | 127 | 10,**24**,25,32,33 | **2.37** | 5 30 0 |
|  | Cingulate Gyrus | 147 | 23,24,31,32 | 2.18 | 5 15 30 |
| **Frontal** | Orbital Gyrus | 29 | 11,47 | 2.31 | 10 55 -20 |
|  | Rectal Gyrus | 44 | 11 | 2.30 | 5 55 -25 |
|  | Inferior Frontal Gyrus | 106 | 9,10,11,13,44,45,46,47 | 2.25 | 10 40 -20 |
|  | Superior Frontal Gyrus | 204 | 8,9,10,11 | 2.33 | 20 50 0 |
|  | Middle Frontal Gyrus | 167 | 6,8,9,10,11,46,47 | 2.31 | 25 55 -5 |
|  | Medial Frontal Gyrus | 243 | 6,8,9,10,11,25 | 2.35 | 10 40 -5 |
|  | Subcallosal Gyrus | 21 | 11,13,25,34,47 | 2.27 | 5 20 -15 |
|  | Precentral Gyrus | 57 | 4,6,9 | 2.07 | -20 -25 55 |
|  | Cingulate Gyrus | 8 | 6,32 | 2.03 | -10 10 40 |
|  | Paracentral Lobule | 46 | 3,4,5,6,31 | 2.09 | -5 -25 45 |
| **Sub-lobar** | Insula | 27 | 13,47 | 2.18 | 30 15 15 |
| **Parietal** | Postcentral Gyrus | 49 | 1,2,3,5,40 | 2.08 | -20 -30 50 |

**Figure 4.**
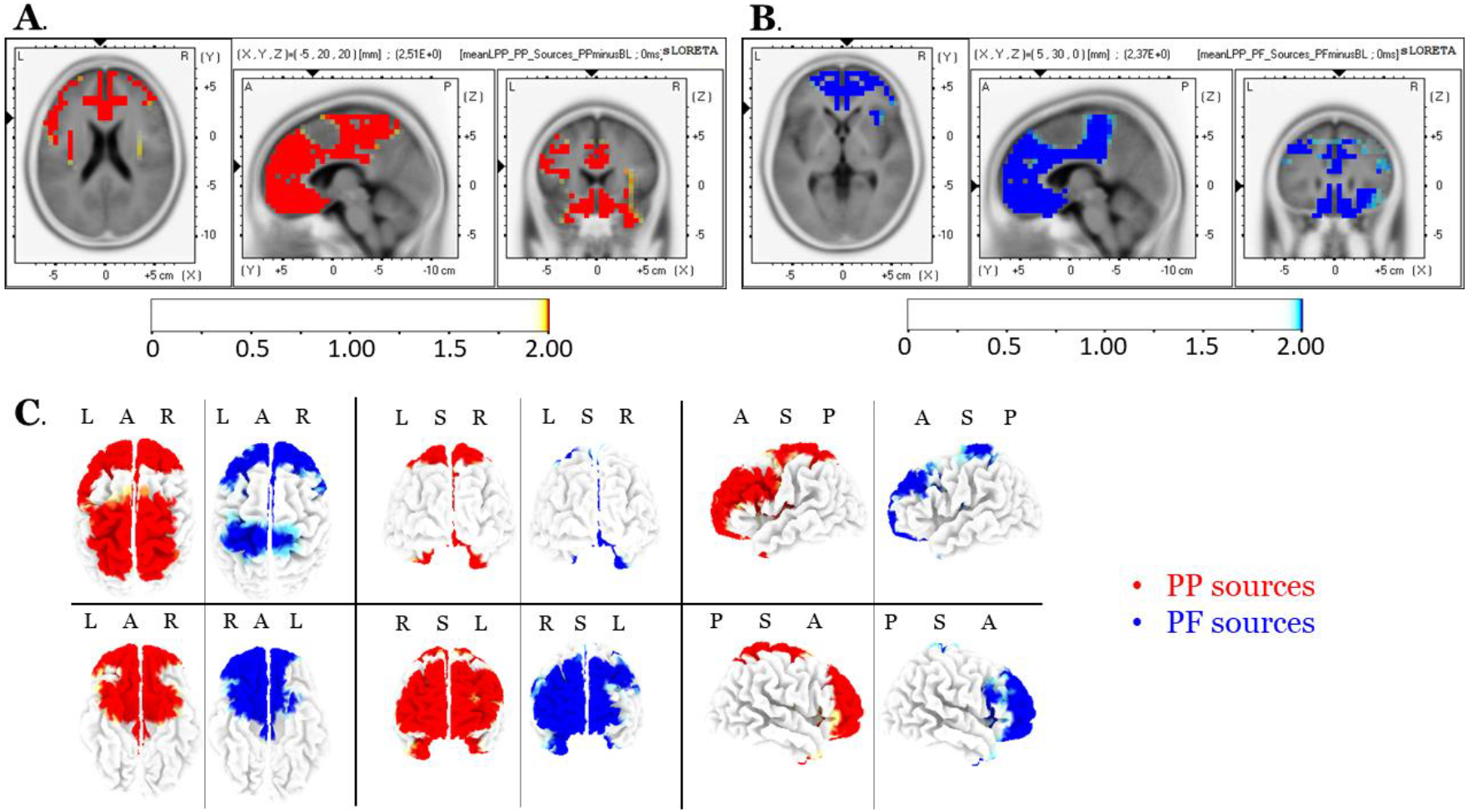
**A.** LPP LORETA slices for maximum-t views associated with the PP sources. **B.** LPP LORETA slices for maximum-t views associated with the PF sources. **C.** Six-view LPP sources in PP (red) and PF (blue) conditions.

There were no significant voxels (all *p*’s>.05) in the PP versus the PF comparisons.

### Band-specific LORETA Sources

We found only alpha-related source differences, with the PP activation greater than that of the PF, predominantly over the right hemisphere central brain regions (specifically, the pre-central gyrus). No other band exhibited significant differences (all *p*>.05). Results of alpha sources are shown in **Fig. 5** and tabulated in **Table 3** (clusters with at least 2 voxels are reported).

**Figure 5.**
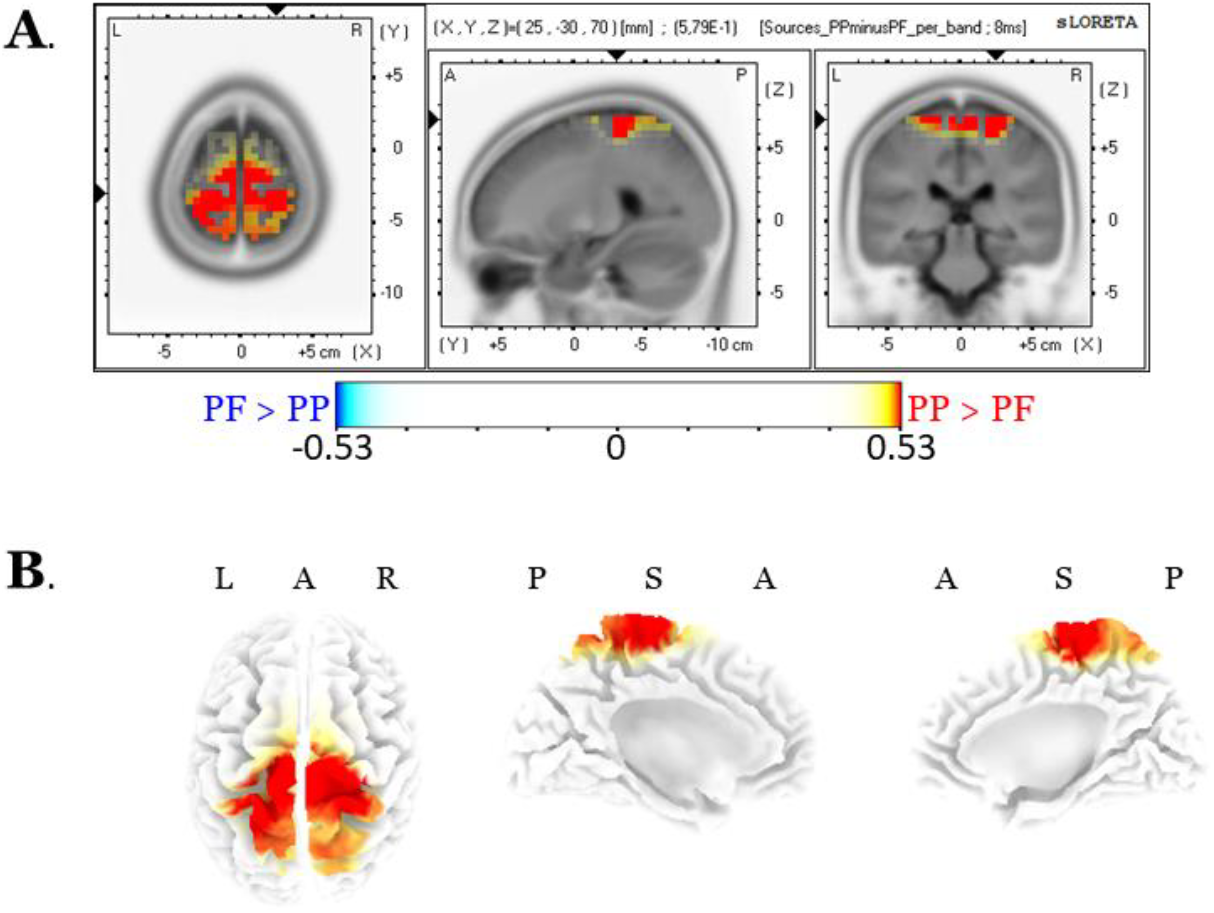
LORETA images of differences in alpha band as (A) xyz-slices and (B) three brain views.

**Table 3.**
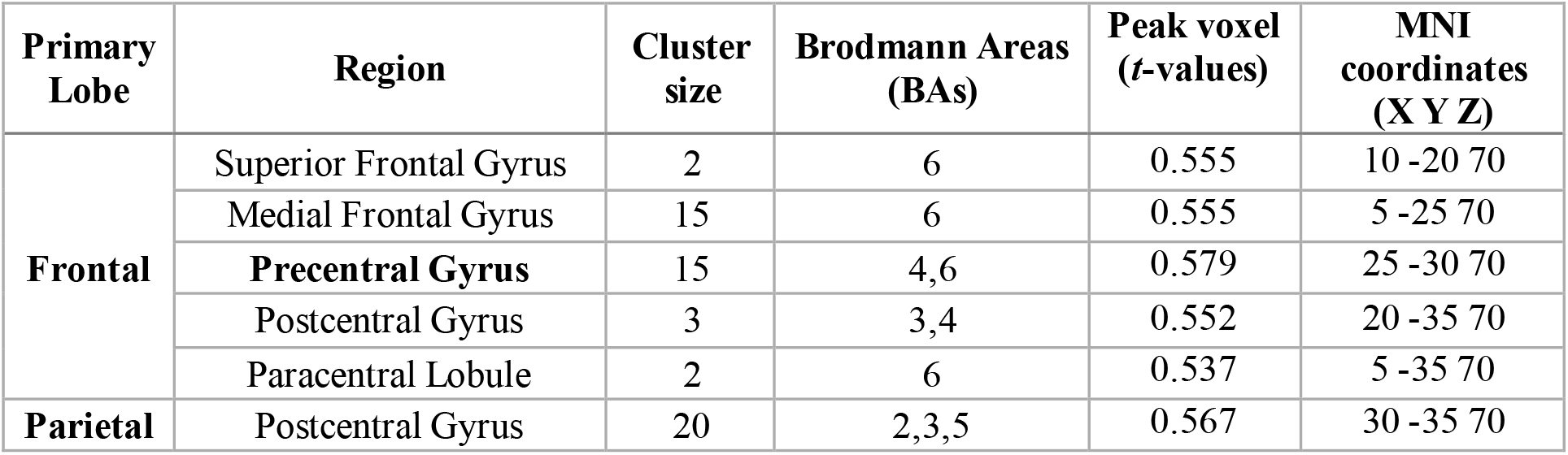
LPP alpha sources that showed higher activations in the PP compared to the PF condition.

| Primary Lobe | Region | Cluster size | Brodmann Areas (BAs) | Peak voxel (t-values) | MNI coordinates (X Y Z) |
| --- | --- | --- | --- | --- | --- |
| Frontal | Superior Frontal Gyrus | 2 | 6 | 0.555 | 10 -20 70 |
|  | Medial Frontal Gyrus | 15 | 6 | 0.555 | 5 -25 70 |
|  | **Precentral Gyrus** | 15 | 4,6 | 0.579 | 25 -30 70 |
|  | Postcentral Gyrus | 3 | 3,4 | 0.552 | 20 -35 70 |
|  | Paracentral Lobule | 2 | 6 | 0.537 | 5 -35 70 |
| Parietal | Postcentral Gyrus | 20 | 2,3,5 | 0.567 | 30 -35 70 |

## DISCUSSION

In the present study, we investigated the brain activity evoked by two mental function s: mental self-projection into the present-past (PP) border, and into the present-future (PF) border. For this purpose, we analyzed the late positive potentials in frontal areas and their cortical generators using the standardized LORETA algorithm. Also, we compared frequency-specific LORETA sources between PP and PF tasks.

We found that the amplitudes of the LPP elicited during self-projection into the PF border were significantly higher than those associated with PP at the left lateral prefrontal areas; interestingly, the opposite patterns were observed in the central and right prefrontal areas. Crucially, for both self-projections (i.e., towards PP and PF borders) the underlying neuronal sources - as reflected by the magnitude of the current densities of the sLORETA vectors - overlapped with the brain’s default mode network and the related interacting areas. Finally, there was enhanced alpha-related activation with respect to PP in comparison to PF, predominantly over the right hemisphere central brain regions (especially over the pre-central gyrus).

To better understand these results, it is beneficial to consider the psychophysiological importance of the alpha EEG oscillations, the genesis of the LPPs, and the procedure used in the sLORETA technique to identify LPP sources (Gallagher, 2019; Nierhaus et al., 2009; Rutiku et al., 2016; Yuan et al., 2009). Moreover, it is useful to consider how the second law of the thermodynamics regarding entropy may apply to these biological systems (Ghaderi, 2019; Parrondo et al., 2009).

Our scalp-domain analyses support the hypothesis that LPPs reflect successful decision making or retrieval during time estimation (19). The observed patterns of LPP alterations are consistent with well-known evidence that the lateral prefrontal cortex is critically involved in temporal order (Fuster,2015; Fuster and Bressler, 2012; Vogeley and Kupke, 2007).

A possible explanation regarding the dissociation of the LPP patterns might be based on the ROBBIA (ROtman-Baycrest Battery to Investigate Attention) model of executive function (Shallice et al., 2008b, 2008a; Stuss, 2011; Stuss and Alexander, 2007); this model postulates that the left–right prefrontal specialization is not only domain-based but also process-based. In particular, the ROBBIA model proposes a prefrontal hemispheric specialization for two distinct executive functions: first, the left-lateralized criterion-setting (or task-setting), which can be defined as the phasic, transient cognitive control processes needed to form or select task-relevant rules (Stuss and Alexander, 2007), suppressing at the same time the task-irrelevant criteria and operations (Vallesi, 2012); and second, the right-lateralized monitoring, which can be defined as the tonic, sustained cognitive control processes needed to actively maintain abstract coded representations of events, monitoring their relative status in relation to each other and their consistency with the intended plan for behavioral adjustments (Petrides, 2006; Stuss and Alexander, 2007; Vallesi, 2012). This phasic-tonic description appears to correspond to potentially alterable (future) versus unaltered (past) events (Callender, 2010; McCormack, 2014; Oppenheim, 2010).

LPP patterns associated with self-projection towards the PP borders are consistent with broad functional and neuroanatomical organizing principles. Such principles indicate that the rostro-caudal axis of the frontal lobes is organized hierarchically, whereby: the posterior PFC areas support control, involving temporally proximate, concrete action representations; conversely, the anterior PFC areas support control, involving temporally extended, abstract representations (Badre,2008; Koechlin and Summerfield, 2007; Petrides, 2006; Wood and Grafman, 2003). Corroborating this notion, lesion studies (Kagerer et al., 2002; Koch et al., 2002), transcranial magnetic stimulation (TMS) (Alexander et al., 2005; Jones et al., 2004; Vallesi et al., 2007), and functional neuroimaging studies (Bueti et al., 2008b, 2008a; Ferrandez et al., 2003; Pouthas et al., 2005) have all implicated the right DLPFC is in time perception.

Statistical comparisons, based on the LORETA analysis, yielded significance thresholds in several brain areas; these thresholds overlapped with the so-called default-mode network (DMN). The DMN includes a set of brain regions, including the medial prefrontal cortex, the lateral and medial parietal cortex (precuneus and retrosplenial cortex), and the lateral and medial temporal lobes, as well as the hippocampus (Mason et al., 2007; Raichle, 2015). It has been extensively documented that the core brain network associated with past and future thinking, operates by engaging several brain regions (including medial prefrontal regions, the posterior regions in the medial and lateral parietal cortex, the precuneus, the retrosplenial cortex, the lateral temporal cortex and the medial temporal lobe). Functionally, past and future thinking involves several cognitive processes, such as episodic thinking, episodic foresight, and related forms of mental construction and simulation. Simulation of *future* events requires the engagement of an unfamiliar (future), rather than a familiar (past) setting, so it differs in the construction and elaboration phases. However, there is convergent evidence that past and future simulation share common brain resources and systems (Bertossi et al., 2016; Lavallee and Persinger,2010; Viard et al., 2011).

The observed alpha-related LPPs differences are consistent with studies supporting the hypothesis that the activation of the prefrontal and premotor areas provides the key mechanism involved in selective attention to time (Nobre and O’Reilly, 2004). Nevertheless,as Milz et al. (2017) reported recently, the enhanced alpha sources during the self-projection into PP borders may reflect a decreased cortical excitability. Specifically, the above authors, by analyzing two 64-channel resting state EEG datasets from healthy participants via exact low-resolution electromagnetic tomography (eLORETA), found that intra-cortical alpha source oscillations reflect decreased cortical excitability. This might be in line with the notion mentioned earlier that ‘the future is open and past is closed’, meaning that events that have not occurred are potentially alterable, unlike the past which is fixed; thus, latter events may be expected to engage lower excitability (Frey et al., 2015; Palva and Palva, 2007).

In accordance with a biological realization of the second law of thermodynamics (namely, that entropy tends to increase), we hypothesized that self-projection into the PF borders would show greater activations than those for PP borders. In other words, considering the stability of the past in contrast to the uncertainty of the future, we expected that the self-projection towards the PF borders would produce enhanced mental effort (higher activations). Extending further the above physical assertions regarding the notion of time, the second law of thermodynamics suggests that, in general, entropy increases over time. In a more precise terminology, entropy defines the extent to which a signal is temporally ordered (low entropy) or unpredictable (high entropy) (Parrondo et al., 2009).

To discuss some of our results we need to introduce the notion of *transfer entropy*. This term, coined by Schreiber (2000), measures the reduction of the uncertainty in inferring the future state of a process, that occurs as a result of the knowledge of the (current and past) states of another similar process. Interestingly, Paluš has shown that transfer entropy can be rewritten as a conditional mutual information (Hlaváčková-Schindler et al., 2007; Paluš et al., 2001). In this sense, although Newton’s time apparently flows equitably, biological time has cyclicity and eddies (Killeen,2014). For example, *déjà vu* entendre throws us back in time, whereas fantasy, planning, and prospective memory throw us forward. In other words, living organisms, by using energy, can decrease, locally, the entropy associated with various cognitive processes. This ability to defy entropy, to extract order from chaos, to predict the future of regular systems, to anticipate prey, and to evade predators, is part of the evolutionary process that allows us to associate events with their specific occurrence in time, independently of past or future (Killeen, 2014).

## CONCLUSION

In conclusion, brain patterns generated by two types of MTT (as reflected by the LORETA technique) showed the activation of a common neural network. Although the two types of self-projection share common cortical resources, they recruit different brain regions in qualitatively different ways; thus, they are associated with specific variations within dissociable large-scale neuroanatomical brain circuits. In this framework, the frontally-distributed LPPs provide a dissociative tool, to elucidate the alternations within the prefrontal cortex that appear during self-projection into the PF vs PP borders, while engaging in WM operations. Finally, considering the notion of thermodynamics, which asserts that entropy tends to increase over time, we did not find fixated patterns of cortical activation in association with self-projection towards the PF borders, despite the expected stability of the past as opposed to the uncertainty of the future. Instead, we found a rather balanced distribution of activation, consistent with the notion that biological time tends to behave in a homeostatic way. The latter suggests that, as a result of appropriate evolutionary processes, we have the capacity to associate events according to their occurrence in time, to extract order from chaos, to predict the future of regular systems, to anticipate prey,and to evade predators (Killeen, 2014).

## ACKNOWLEDGEMENTS

The authors warmly thank Dr. Alexandros Pino (Ph.D., Laboratory Teaching Staff at National and Kapodistrian University of Athens, Greece) and Emmanouil A. Kitsonas (Electrical Engineer, Ph.D. Technical Director, Eugenides Foundation Member TCG, IEEE, IPS Council) for their support regarding the construction of the experimental procedure. The authors also thank Dr. Antonis Mavromatos (University Mental Health, Neurosciences and Precision Medicine Research Institute “COSTAS STEFANIS”, (UMHRI), Athens, Greece) for his contributions to the experimental evaluation.

## AUTHOR CONTRIBUTIONS

Conceptualization, C.P.; Methodology, C.P., A.E.G., A.S.F., and C.N.C.; Software, A.E.G., N.C.K., and P.P.; Validation, C.P., A.S.F., P.M.T. and C.N.C.; Formal Analysis, A.E.G., N.C.K., and P.P.; Investigation, C.P., and A.S.F.; Resources, C.P., P.P., and X.S.; Data Curation, C.P. and X.S.; Writing – Original Draft, C.P., A.E.G., A.S.F., and P.M.T.; Writing – Review and Editing, C.P., A.S.F. and P.M.T..; Visualization, A.E.G., N.C.K., and P.P.; All authors reviewed the paper and approved the submitted version.

## ETHICS DECLARATIONS

The study was approved by the University Mental Health, Neurosciences and Precision Medicine Research Institute “COSTAS STEFANIS” (UMHRI). All procedures performed in “Protagoras” experiment were in accordance with the ethical standards of the institutional and/or national research committee and with the 1964 Helsinki declaration and its later amendments or comparable ethical standards for experiments involving human participants. Informed consent was obtained from all individual participants included in the study.

## DATA AND CODE AVAILABILITY

Data and codes (software application and custom scripts) supporting the study findings are available from the authors upon request.

## APPENDIX Why do we use the term “Borders of Present”?

With some degree of abstraction, we use the term “borders (or limits) of the present”, in analogy with the use of this term in mathematics. The terms “past” and “future” are used in their traditional form: the past refers to those events that occurred before a given point in time. The future is the portion of the projected time direction that is anticipated to occur; depending on the context, it may be have an infinite extent, or it may be circumscribed and finite. The definition of the term “present” is a slightly more abstract: it may be defined as the time associated with events perceived directly and for the first time, i.e., it is not considered as a recollection of the past or as a speculation of the future. It is equivalent to the word “now”, and is the period of time located between the past and the future.

At this point, it is worth pointing out a fundamental mathematical notion arising in number theory. According to ancient philosophical considerations, the physical number “One” is the fundamental “unit” that generates all numbers: all other real numbers can be generated by applying repeatedly additions and multiplications starting with the number “One”. Actually, (1 + *ε*)^*n*^ = *real number* (*rational or irrational*) But what happens if we add some elemental quantity to“One” and keep on repeating this process at *infinitum*? In mathematical terms, this question takes the form of computing the limit, 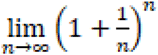. This limit gives rise to the so-called Euler’s number (*e* ≅ 2.72), which is one of the most important (irrational) numbers in mathematics.

Considering the above notions, as well as the philosophical position that our existence might be conceived as the propagation (or proliferation) of our present (‘par-on’ in Greek, i.e., close to being), then it follows that it is important tounderstandthe cognitive representations of the borders (or limits) of the concepts “present in relation to both “past” and “future”. Associating concepts arising in MTT with analogous mathematical notions, the number “One” could perhaps be associated with the starting-point (or “unit”) of MTT, namely with the “present”. This unit is projected from “present” to“near past” or “near future” (i.e., 1 ± *ε*); then, it is propagated forwards or backwards (i.e. 1 ± *ε*)^*n*^), resulting, respectively, in a prediction or a memory.

The present study adopted a theoretical strategy by using the verb tenses (Past, Present, Future) as an entree to conceptual representations relevant to time. By analyzing the electrophysiological activity associated with these representations, attempted to elucidate some of the neurological mechanisms underlying this challenging area. In particular, it addressed the following important question: Are the fundamental types of self-projection relatedtotime, namely PP and PF borders, identical with respec t to the elicited LPP responses?

